# A dynamic epigenetic perspective on above and below-ground phenotypic responses to drought: insights from global DNA methylation in *Erodium cicutarium*

**DOI:** 10.1101/2024.04.01.587556

**Authors:** Conchita Alonso, Mónica Medrano, Carlos M. Herrera

## Abstract

- **Premise of the study**. Mounting evidence supports the view that the responses of plants to environmental stress are mediated by epigenetic factors, including DNA methylation. Understanding the relationships between DNA methylation, plant development and individual fitness under contrasting environments is key to uncover the potential impact of epigenetic regulation on plant adaptation. Experimental approaches that combine a controlled alteration of epigenetic features with exposure to some relevant stress factor can contribute to this end.
- **Methods**. We combined the experimental application of a demethylating agent (5-azacytidine) with recurrent drought, and recorded their effects on above- and below-ground phenotypic traits related to early development, phenology and fitness in *Erodium cicutarium* from two provenances.
- **Key results**. We found that 5-azacytidine significantly reduced DNA methylation in leaf and root tissues. Moreover, it slowed plant development, delayed flowering, and reduced the number of inflorescences produced, and such detrimental effects occurred independently of water regime. Recurrent drought reduced final above- and below-ground biomass and total inflorescence production, and such negative effects were unaffected by artificial changes in DNA methylation. Increased fruit and seed-set were the only adaptive responses to drought observed in *E. cicutarium*, together with an increased number of flowers per inflorescence recorded in water stressed plants previously treated with 5-azacytidine.
- **Conclusion**. Epigenetic effects can desynchronize plant growth, flowering and senescence among individual plants in both favourable and adverse environments. Future studies should focus on understanding intraspecific variation in the ability to change plant methylome in response to stress.

## Introduction

Understanding how plant phenotype and fitness change under stressful conditions is a fundamental question in evolutionary ecology with potential application in agronomy and biodiversity conservation. The traditional elements considered to explain observed phenotypic variation include the genotype, which refers to any type of genetic grouping such as clones, siblings, varieties or populations, the environment (i.e., the level of the stress), and the genotype-by-environment interaction that evaluates whether certain genotypes respond differently to environmental variation (Bradshaw, 1965; Schlichting, 1986). More recently, studies have revealed that epigenetic factors such as DNA methylation, histone modifications and small RNAs, participate also in plant phenotypic variation and response to stress (e.g., Chinnusamy and Zhu, 2009; Mirouze and Paszkowski, 2011; Gutzat & Mittelsten Scheid, 2012; Rodríguez López and Wilkinson, 2015; Banerjee and Roychoudhury, 2017), being further related to phenotypic plasticity, i.e., the ability of a certain genotype to produce different phenotypes in different environments (e.g., Herman *et al*., 2014; Kooke *et al.,* 2015; Schneider, 2022). Epigenetic variants (in contrast to conventional genetic mutations) are reversible, more frequent, and they can be to some extent induced by the environment (Lloyd and Lister, 2022), favoring that adaptive adjustments appear in several individuals in a population (Burggren, 2016). Altogether, at least in part, epigenetics could explain how phenotypic plasticity arises quickly and could evolve under different environmental scenarios (Jablonka, 2013; Zhang *et al.,* 2013; Herman *et al.,* 2014; Kooke *et al.,* 2015; Burggren, 2016).

In particular, drought is one of the major environmental stress factors that can limit seriously plant growth and productivity. Different plant species have evolved distinct strategies in order to reduce its deleterious effects depending on the extent of water deficit and its life-history features (Chaves *et al.,* 2002; Kooyers, 2015; Marin de la Rosa *et al.,* 2019). The genetic basis for phenotypic responses to drought, such as changes in root growth and water use efficiency, together with extensive genetic variation in drought resistance into natural populations have been documented in some species (reviewed by Kooyers, 2015). Furthermore, epigenetic changes after drought stress have been frequently observed in experimental settings with crop and model plant species (e.g., Bossdorf *et al*., 2010; Bertolini *et al.,* 2013; Zheng *et al.,* 2017; Neves *et al.,* 2017) although the consequences for individual fitness are infrequently reported (but see Zhang *et al.,* 2013; VanDooren *et al.,* 2020). Experimental modification of DNA cytosine methylation offers a suitable approach to evaluate the impact of this epigenetic mechanism on phenotypic traits, plant development and final individual fitness expressed under contrasting environmental conditions (see e.g., Herman and Sultan, 2016; Rendina, *et al.,* 2016; Münzbergová *et al*., 2019). The use of inhibitors of the activity of DNA methyltransferase enzymes such as 5-azacytidine, 5-aza-2’-deoxycytidine or zebularine provides a tractable way to modify DNA cytosine methylation in plants, which could enlarge natural variation in this particular epigenetic feature, while controlling for genetic relatedness (Alonso *et al.,* 2017; Puy *et al.,* 2017; Herrera *et al.,* 2019).

Spatially heterogeneous and unpredictable environments are expected to favor phenotypic plasticity and, thus, epigenetic responses to stress, because the direction and strength of selection forces are variable in space and/or time (Bradshaw, 1965; Hendry, 2016). Mediterranean mountains provide a good example of heterogeneous and unpredictable environments associated to seasonal and interannual rainfall variation and topographic complexity (see e.g., Cowling *et al.,* 2005; Ramos-Calzado *et al.,* 2008; Cook *et al.,* 2016). Annuals frequently adopt there a scape-avoidance strategy that can be expressed through escape (i.e., early flowering to complete life-cycle before drought becomes severe) and/or changes in root growth and water use efficiency to reduce fitness costs (Chaves *et al.,* 2002; Franks, 2011; Kooyers, 2015). In this study, we investigated the potential role of DNA methylation in determining the phenotypic responses of *Erodium cicutarium* (Geraniaceae) to recurrent drought. We applied 5-azacytidine at seed germination to experimentally alter DNA cytosine methylation of sibling individuals and grew them under two different water regimes, following a factorial design. As *E. cicutarium* is a fast growing annual that flowers in spring, when precipitation is largely unpredictable (Cowling *et al.,* 2005), we applied a recurrent drought treatment that started after a period for individuals’ establishment. Recurrent stress with recovery periods is thought to elicit a distinct and stronger epigenomic signal than single extreme events and it is expected to promote stress memory and tolerance to subsequent stress (Walters *et al.,* 2011; Fleta-Soriano and Munné-Bosch, 2016). We first checked whether the demethylation treatment, which effectively reduced cytosine methylation at the seedling stage in this species (Alonso *et al.,* 2017), actually remained detectable in leaf and root tissues at the adult stage. Further, we explored the consequences of seed demethylation and recurrent drought in terms of plant development (first leaf length, time to first flower, changes in leaf and inflorescence numbers over time) and final (static) phenotype (reproductive output, above and below-ground biomass). We tested the following predictions: (1) Reduction of DNA methylation at seed germination will impact on subsequent plant development, phenology and fitness. (2) Recurrent drought will induce a plastic response in terms of plant development, phenology and/or fitness. (3) If DNA methylation is involved in drought-induced plasticity, then demethylation should blur or impair the phenotypic responses to drought.

## Materials and Methods

### Study species and experimental design

*Erodium cicutarium* (L.) L’Hér. (Geraniaceae), is an annual herb native to Mediterranean Europe, North Africa and western Asia, that is currently distributed globally in temperate areas with hot summers of both hemispheres (Fiz-Palacios *et al.,* 2010). A fast growing cycle, which allows early flowering and fruiting before seasonal summer drought, and an autonomous pollination system have together contributed to its naturalization in diverse habitats from all continents (see e.g., Francis *et al.,* 2012; Kimball *et al.,* 2014).

In this study, we analyzed the effects of the demethylating agent 5-azacytidine and recurrent drought on plant phenotypic traits related to early development, phenology, growth, and fitness, and the level of global DNA cytosine methylation in the genomes of their leaf and root tissues. Plants were grown in the greenhouse for two generations to minimize the effects that heterogeneous natural growing conditions of the maternal plants might induce in the epigenetic response to changes in specific environmental factors of their offspring (Latzel, 2015).

In spring 2015, we collected mature fruits from adult plants at two *E. cicutarium* natural populations located in Cazorla mountains (Jaén province, SE Spain): Puerto del Tejo (PT, 1590 m asl) and Nava de las Correhuelas (CH, 1625 m asl) distant ca. 10 km in straight-line. Puerto del Tejo population has a steeper slope and more rocky substrate, and individuals at this site were larger in diameter but produced fewer leaves than those at CH population (Alonso *et al.,* 2017). Fruits were stored in darkness at room temperature for 4-5 months. In autumn 2015, seeds were removed from fruits, scarified and germinated in universal substrate (COMPO SANA®) mixed in 3:1 with perlite (“substrate” hereafter). These first generation (F1) seedlings were subsequently transferred to 1L pots with the same substrate; pots were grouped in trays that were periodically rotated within the greenhouse (16h light; 25-20 °C) until the end of reproduction (ca. 6 months). Half of the trays were watered twice per week and the other half were watered once every 10-11 days (see below for details). Autonomously-pollinated fruits were collected in paper bags, and stored at room temperature.

In October 2016, seeds from six mothers from each of the two water regimes and population (N = 24 families: 2 populations * 2 water maternal regimes * 6 F1-mothers) were weighed and slightly scarified with sandpaper. A demethylation treatment was applied to half of the seeds as described in Alonso *et al*. (2017). In brief, scarified seeds were submerged in 150 µl of either Control (water with DMSO 97:3, v:v) or a 0.5 mM solution of 5-azacytidine (Sigma A2385-100mg; 5azaC hereafter) during 48 h at 4 °C. All seeds were then individually transferred to seedling plug trays filled up with substrate. Trays were saturated in water and placed under greenhouse conditions (16h light; 25-20 °C). Twenty days after sowing, 288 seedlings (144 control and 144 treated with 5azaC) of these second generation (F2) were transplanted into 1L pots. Other F2 extra seedlings (N = 40) were collected a few days later and processed to confirm the efficiency of the demethylation treatment at this stage (see results in Alonso *et al.,* 2017).

The recurrent drought treatment started three weeks after transplant, when seedlings were around six weeks old. For each maternal line (line, hereafter), half the offspring was watered at field capacity twice per week and half the offspring was watered at field capacity once every 10-11 d until the end of the experiment. The full design was a 2 x 2 factorial with three replicates per line. Flowering started one week after transplant and reached the peak (75 % individuals with flowers) at week 10. The experiment finished when plants were 17 weeks old and plants started showing signs of senescence.

### Trait data collection

For each F2 trasplanted plant (N = 288), we recorded developmental traits at different times (ontogenic approach) and final fitness-related traits (static approach). The ontogenic -or longitudinal-approach aimed to characterize initial size (first leaf length) and changes associated to growth and phenology by measuring maximum leaf length before transplant and periodically monitoring flowering (e.g., time to first flower) and recording the number of leaves and inflorescences plants had. The number of full-grown leaves was counted twice before recurrent drought started, when plants were 11 days and ca. five weeks old. Leaf counting was repeated at week nine, after a month of the onset of drought treatment. Inflorescences were first counted at week nine, when still more than 60 % of individuals remained vegetative. Inflorescence counting was repeated at week 13, when 99 % plants had flowered, and finally at week 17, when 14 % plants showed signs of senescence. In all cases, only full-grown non-senescent leaves were recorded and the petiole of counted inflorescences was marked to be sure they were not counted twice.

The static approach looked at reproductive fitness (number of total inflorescences produced, number of flowers per inflorescence, proportion of flowers that set fruit, proportion of seeds per ovule) and final individual size (above- and below-ground biomass). For each plant, above-ground biomass was collected at the end of the experiment, placed in labeled individual paper bags and dried at 50 °C for at least 48 h before further processing. For a subsample (N = 96 plants), roots were taken out of soil, carefully washed in water, excess water was wiped away using absorbent paper and a sample of fine roots (avoiding the primary thickest one) was collected for DNA analyses, placed in a labeled paper bag and dried at ambient temperature in sealed containers with abundant silica gel. The rest of the root was placed in labeled paper bags and dried at 50 °C for a week before further processing. Just before weighing, the bags were placed again in the oven for > 8 h, inflorescences were separated from the vegetative tissues and each part was weighed in a digital balance (Mettler-Toledo XS-105) to the nearest 0.001 g. Biomass of reproductive and vegetative tissues were positively correlated across samples (rs = 0.59, P < 0.0001, N = 212). Thus, we will present only the analyses regarding total above-ground mass because they were consistent with those observed for the two types of tissues. The same protocol was followed for weighing the roots. Reproductive success was estimated as the proportion of flowers setting fruits (hereafter named fruit-set) and the proportion of ovules setting seeds (seed-set, hereafter), and for each parameter an average value per plant was computed on a per inflorescence basis. In both cases, the variable was far from a normal distribution because a large number of individuals had a 100 % of success with median value for each proportion being equal to 0.91 for fruit-set and 0.83 for seed-set.

### Sample processing and laboratory methods

Once each F2 individual reached the flowering stage, a sample of 2-3 full grown leaves without signs of damage or senescence of each individual was collected, placed in labeled paper bags and dried at ambient temperature in sealed containers with abundant silica gel. Each root sample collected at the end of the experiment was handled similarly as mentioned above.

Dried leaf and root tissues were homogenized to a fine powder using a Retsch MM 200 mill. Total genomic DNA was extracted using Bioline ISOLATE II Plant DNA Kit and quantified using a Qubit fluorometer 2.0 (Thermo Fisher Scientific, Waltham, MA, USA). A 100 ng aliquot of DNA extract was digested with 3 U of DNA Degradase PlusTM (Zymo Research, Irvine, CA, USA), a nuclease mix that degrades DNA to its individual nucleoside components. Digestion was carried out in a 40 µl volume at 37 °C for 3 h, and terminated by heat inactivation at 70 °C for 20 min. Two independent replicates of digested DNA per sample were processed to estimate global cytosine methylation more precisely. Selective derivatization of cytosine moieties with 2-bromoacetophenone under anhydrous conditions and subsequent reverse phase HPLC with spectrofluorimetric detection, were conducted. For each tissue, the 192 vials (96 samples x 2 replicates) were processed in randomized order. One leaf replicate failed to produce any signal and was discarded (N = 191); three additional replicates were run for root samples (N = 195). The percentage of total cytosine methylation on each replicate was estimated as 100*5mdC/(5mdC + dC), where 5mdC and dC are the integrated areas under the peaks for 5-methyl-2’-deoxycytidine and 2’-deoxycytidine, respectively (see Alonso *et al.,* 2016 for further details).

### Statistical analyses

All statistical analyses were carried out using the R environment (R Development Core Team 2019, R version 3.6.3). All data were visually inspected and, for some variables, obvious outliers (N ≤ 3) were excluded on the assumption they were originated from some anomaly or failure in the processing. For those variables that were measured in several modules of each plant (e.g., number of flowers per inflorescence) or estimated twice (e.g., global cytosine methylation percentage) the average value was calculated and individual data used as response variable. For reproductive success variables (i.e., fruit-set and seed-set proportions), which have highly skewed distributions, we used the standardized variable as response.

Linear models were used to analyze the fixed effect of the two experimental factors, namely demethylation (control *vs*. 5azaC) and recurrent drought (watered *vs*. drought), and their interaction on all study traits (except for the early phenotypic traits measured before the recurrent drought where only demethylation was included). The lme function from the nlme package (Pinheiro *et al.,* 2017) was used to fit mixed-effect models with Restricted Maximum Likelihood (REML) estimation, including maternal family as a random effect (Bolker, 2015). The analysis of the possible effect of the maternal water regime was out of the scope of this study, thus it was included as a random grouping factor in which maternal family was nested, this procedure corrected for the non-independence of data and ensured that any possible influence of family heterogeneity in genetic and environment backgrounds were adequately accounted for (i.e., blocked; Mead, 1988). The effect of provenance (PT *vs*. CH) was considered fixed because, although it was not the focus of our research, the two provenances are known to differ in the consequence of demethylation treatment for DNA global methylation of seedling roots (Alonso *et al.,* 2017). When provenance had significant interactions with any of the two experimental factors, partial models for each population were run separately to better understand the effects of the focal experimental factors. In the case of cytosine methylation percentage, the full model included tissue (leaf vs. root), demethylation, recurrent drought, provenance, and all their interactions as fixed effects. Partial models were subsequently applied to leaf and root tissue independently to better interpreting the sign of observed interactions.

Plant developmental traits were characterized by changes with time in the no. leaves and the no. inflorescences per individual. The two variables were analyzed as repeated measurements with lmer function from the lme4 package (Bates *et al.,* 2015), by modelling time as an ordered fixed factor with three levels, including individual as a categorical random factor, and explicitly analyzing the interactions between time and the three experimental factors [i.e., response variable ∼ time*provenance + time*demethylation + time*drought + (1 | individual)].

For every fitted model, inspection of residuals and goodness-of-fit assessment was conducted using the function check_model of Performance package (Lüdecke *et al.,* 2021). Significance of fixed effects and interactions were obtained by applying to fitted models the Anova function from the car package (Fox and Weisberg 2011). Model-adjusted marginal averages for the fixed main effects were obtained with the emmeans function from the emmeans package (Lenth *et al.,* 2019).

## Results

### Global DNA methylation at adult stage

In order to test the efficacy of the seed demethylation treatment, we tested for its specific effect only in those plants that were well-watered along their development (i.e., they did not experience any drought). In this subset, the effect of seed demethylation was highly statistically significant (ꭓ^2^ = 24.6, df = 1, P < 0.0001), similar for the two provenances (ꭓ^2^ = 1.35, df = 1, P = 0.24), and more evident on root than in leaf DNA (ꭓ^2^ = 3.2, df = 1, P = 0.07, for the demethylation*tissue interaction; Fig. 1). We also found that, at this adult stage, DNA cytosines were more frequently methylated in roots than leaves (ꭓ^2^ = 41.9, df = 1, P < 0.0001; 22.8 % ± 0.3 *vs*. 20.9 % ± 0.3, for root and leaf tissue, respectively).

**Figure 1.**
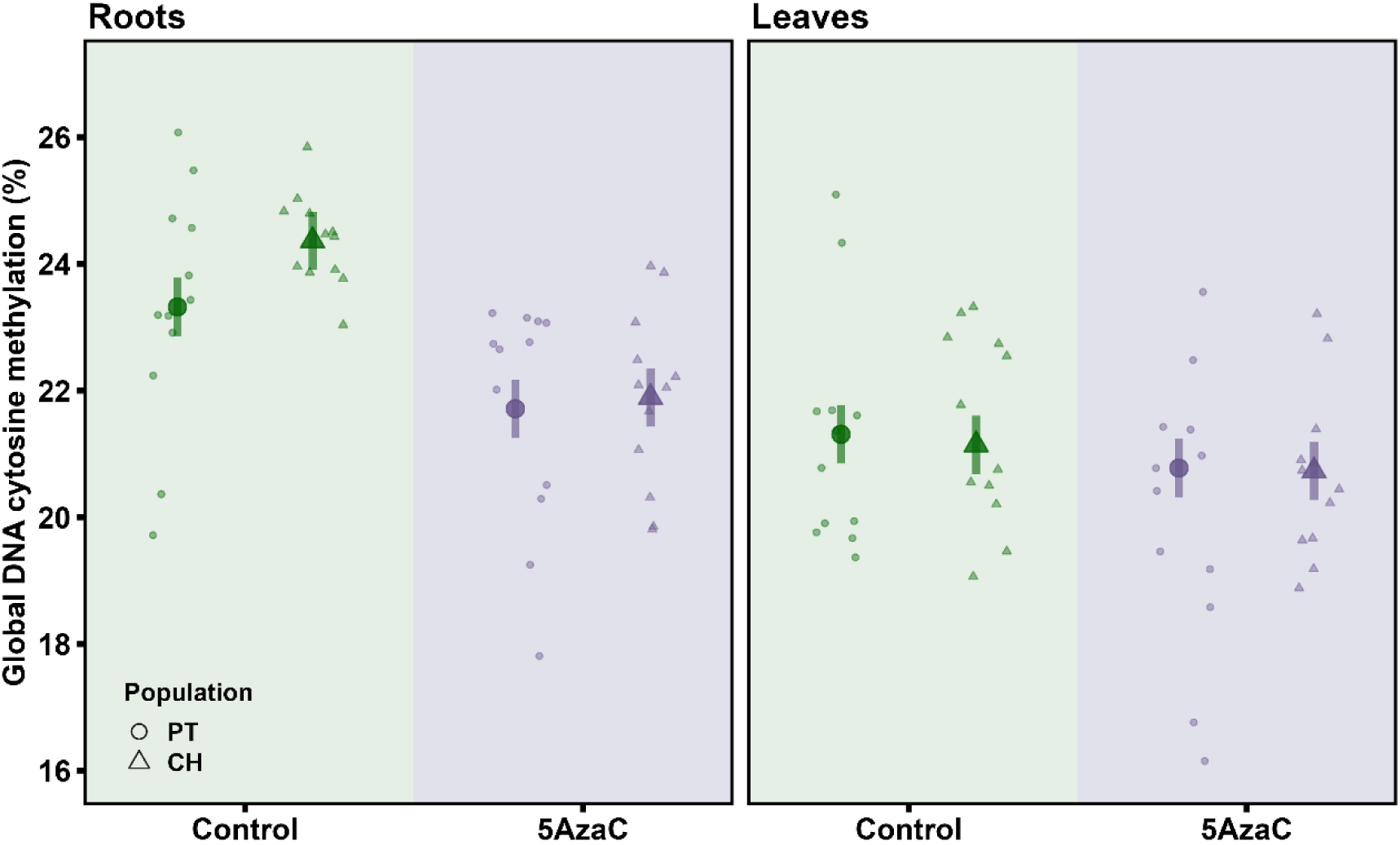
Effects of seed demethylation treatment in global percent of methylated cytosines in DNA obtained from leaves and roots of *Erodium cicutarium* adult plants that did not experience any drought along development. Small circles and triangles denote individuals from PT and CH population, respectively. Big symbols and bars represent leas-squared means and their 95% confidence level, respectively.

In the full experimental design (i.e., including well-watered and drought plants), global DNA methylation in leaves was poorly explained by the fixed factors (Table 1A). In contrast, for root DNA we found multiple significant effects. Both demethylation and drought significantly reduced global DNA methylation, which was also lower in roots of PT plants (Table 1B). We also found significant interactions between drought and provenance and demethylation and provenance, altogether indicating that the effect of drought was stronger in PT plants and, interestingly, the effect of 5azaC was erased in plants experiencing recurrent drought (Table 1B; Supp. Mat. Fig. S1).

**Table 1.**
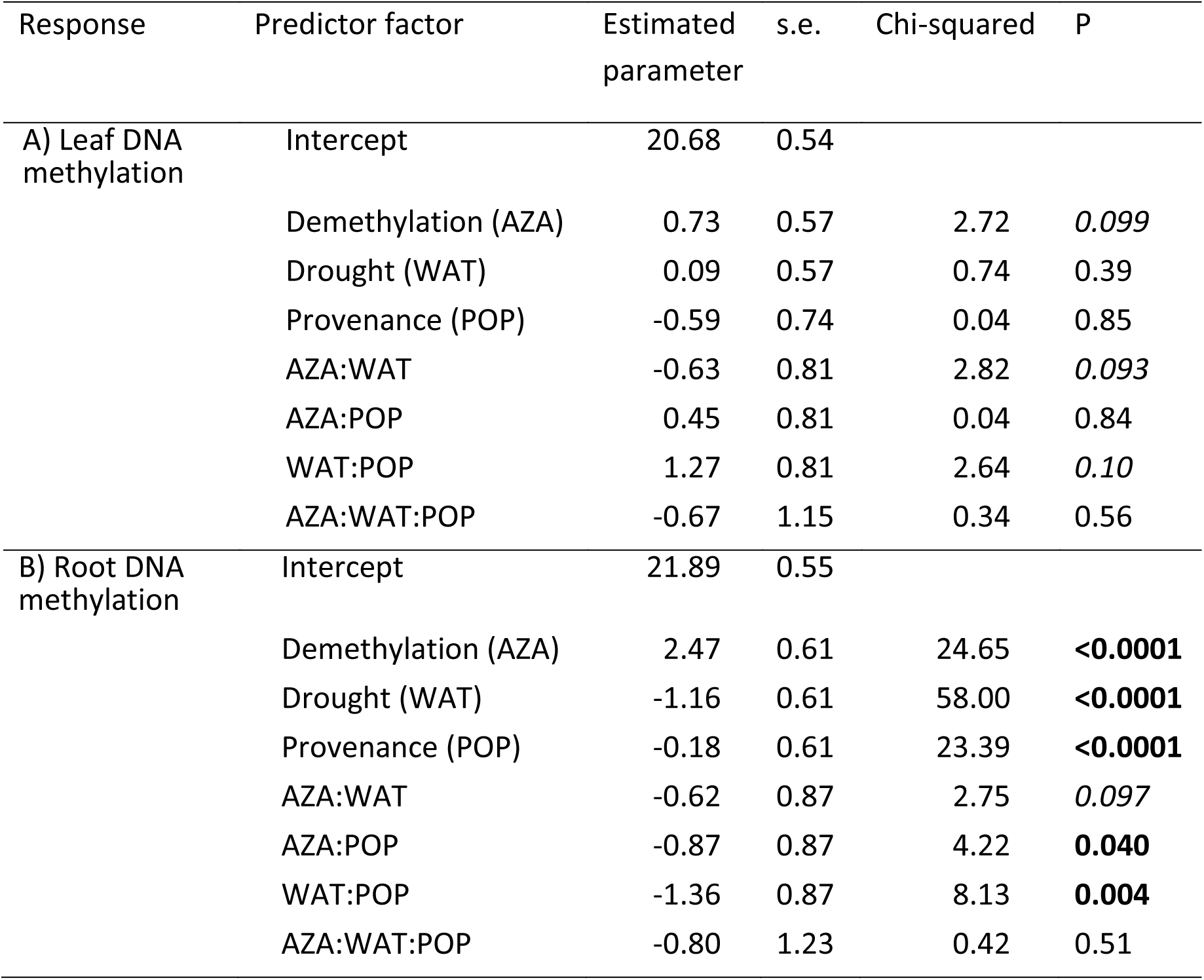
Summary of the results of the linear mixed models conducted to analyze the effects of seed demethylation, water recurrent stress, plant provenance and all interactions on the observed variation in global cytosine DNA methylation percentage of *Erodium cicutarium* adult plants grown in the greenhouse. A) Leaf tissue. B) Root tissue. Each individual plant was represented by the average value of the two to four replicates per tissue analyzed (N = 96 individuals). The significance of the effects are highlighted in bold when P < 0.05 and italics when 0.05 < P ≤ 0.10.

| Response | Predictor factor | Estimated parameter | s.e. | Chi-squared | P |
| --- | --- | --- | --- | --- | --- |
| A) Leaf DNA methylation | Intercept | 20.68 | 0.54 |  |  |
|  | Demethylation (AZA) | 0.73 | 0.57 | 2.72 | *0.099* |
|  | Drought (WAT) | 0.09 | 0.57 | 0.74 | 0.39 |
|  | Provenance (POP) | -0.59 | 0.74 | 0.04 | 0.85 |
|  | AZA:WAT | -0.63 | 0.81 | 2.82 | *0.093* |
|  | AZA:POP | 0.45 | 0.81 | 0.04 | 0.84 |
|  | WAT:POP | 1.27 | 0.81 | 2.64 | *0.10* |
|  | AZA:WAT:POP | -0.67 | 1.15 | 0.34 | 0.56 |
| B) Root DNA methylation | Intercept | 21.89 | 0.55 |  |  |
|  | Demethylation (AZA) | 2.47 | 0.61 | 24.65 | **<0.0001** |
|  | Drought (WAT) | -1.16 | 0.61 | 58.00 | **<0.0001** |
|  | Provenance (POP) | -0.18 | 0.61 | 23.39 | **<0.0001** |
|  | AZA:WAT | -0.62 | 0.87 | 2.75 | *0.097* |
|  | AZA:POP | -0.87 | 0.87 | 4.22 | **0.040** |
|  | WAT:POP | -1.36 | 0.87 | 8.13 | **0.004** |
|  | AZA:WAT:POP | -0.80 | 1.23 | 0.42 | 0.51 |

Moreover, we found that cytosine methylation in DNA obtained from adult leaves and roots was positively related in well-watered plants (r = 0.33, P = 0.02, N = 48). In contrast, in those individuals experiencing recurrent drought, global DNA methylation in the two tissues was not significantly correlated (r = -0.15, P = 0.31, N = 48; Supp. Mat. Fig. S2).

### Phenotypic traits prior to recurrent drought

Germination was very successful in this trial, 42.7 % individuals developed a radicle during the 48 h imbibition period, and 95 % of seedlings showed their cotyledons before 48 h after sowing in soil. The length of the first true leaf was shorter in individuals previously treated with 5azaC (58.1 ± 2.2 mm and 84.0 ± 2.2 mm, for treated and control plants respectively; ꭓ^2^ = 103.3, df = 1, P < 0.0001; Fig. 2A). We also found a significant interaction between demethylation and provenance (ꭓ^2^ = 6.5, df = 1, P = 0.01), the reduction of leaf length after 5azaC treatment being larger in CH plants (Fig. 2A). After 11 d of sowing, seedlings produced between 0 - 8 true leaves, and on average those treated with 5azaC produced fewer leaves than the untreated ones (3.7 ± 0.1 leaves and 4.9 ± 0.1 leaves for treated and control plants respectively; ꭓ^2^ = 51.2, df = 1, P < 0.0001). Again, the effect of 5azaC tended to be stronger in CH plants, as suggested by the marginally significant interaction between demethylation and provenance (ꭓ^2^ = 3.7, df = 1, P = 0.05).

**Figure 2.**
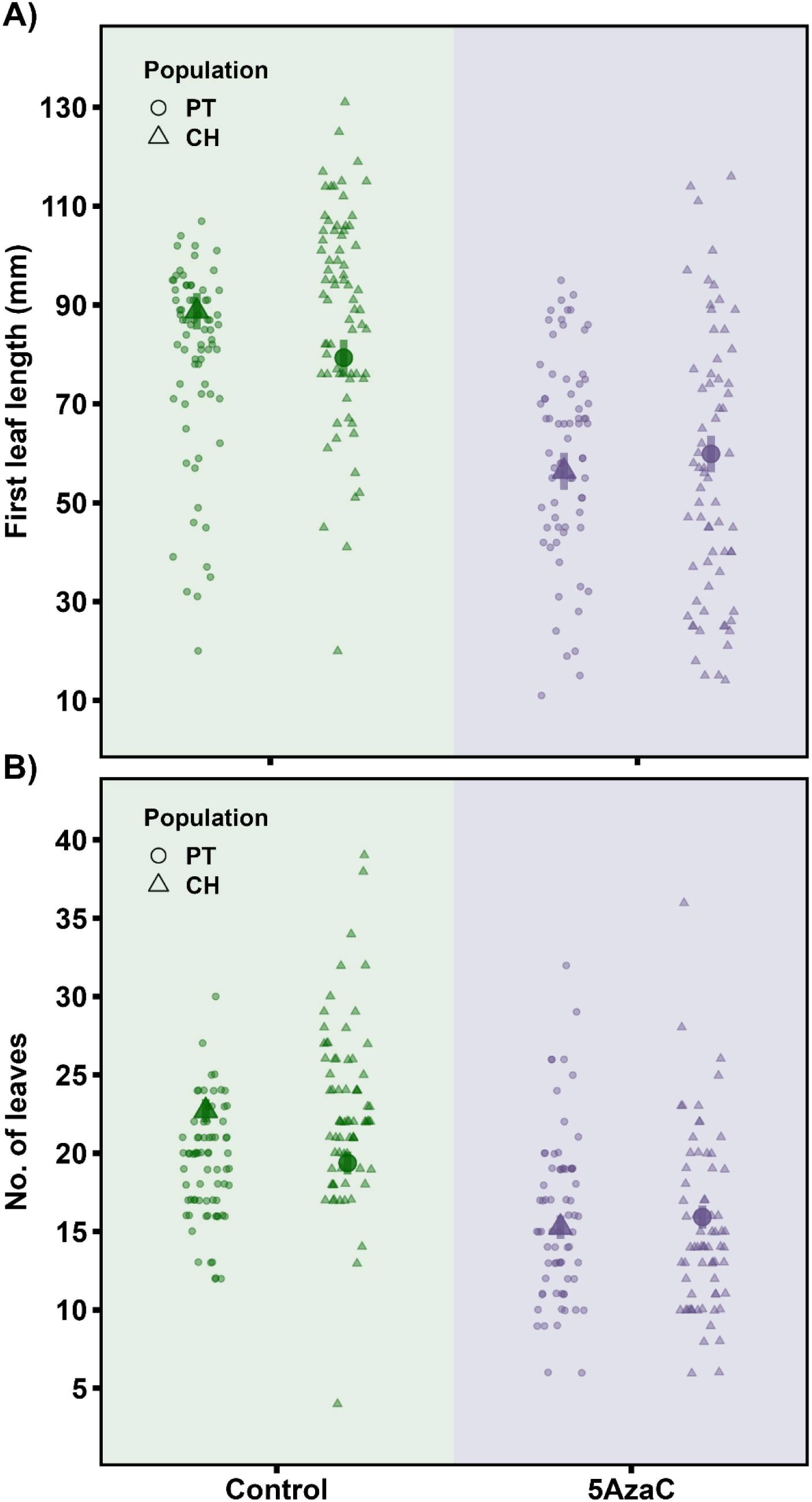
Effects of seed demethylation treatment in early growth traits of *Erodium cicutarium*. A) Length of the first true leaf (mm). B) Number of leaves produced after four weeks of sowing. Small circles and triangles denote individuals from PT and CH population, respectively. Big symbols and bars represent leas-squared means and their 95% confidence level, respectively.

In the four weeks between the first counting and the initiation of the drought treatment, individual plants produced between 2 - 39 leaves. This variance was explained by a significant reduction of leaf production in plants treated with 5azaC, that on average produced 7.4 fewer leaves than their untreated relatives (ꭓ^2^ = 91.9, df = 1, P < 0.0001; Fig. 2B). In addition, a significant interaction between demethylation and provenance indicated that such reduction was stronger in CH plants (ꭓ^2^ = 12.4, df = 1, P = 0.0004; Fig. 2B). The length of the longest leaf at this time ranged between 33 and 279 mm. This variance was explained by the combined effect of demethylation and provenance. First, plants treated with 5azaC had shorter leaves (ꭓ^2^ = 64.05, df = 1, P < 0.0001). Similarly, plants from CH population had leaves than were on average 12.5 mm longer than those from PT population (ꭓ^2^ = 10.9, df = 1, P = 0.0009) and experienced a stronger reduction in length after seed demethylation (ꭓ^2^ = 4.95, df = 1, P = 0.026).

### Developmental traits: flowering phenology and vegetative growth

Both demethylation and provenance had significant effects on the time required to start flowering and no significant interaction between factors was found (Fig. 3). On average, plants from PT started flowering after two months (60.6 ± 1.7 days) whereas those from the CH population started flowering 10.6 days later (ꭓ^2^ = 19.9, df = 1, P < 0.0001; Fig. 3). Further, demethylated plants tend to flower 5.1 ± 1.3 days later than their untreated relatives (ꭓ^2^ = 27.7, df = 1, P < 0.0001; Fig. 3). The effect of recurrent drought was not statistically significant for this trait (ꭓ^2^ = 1.5, df = 1, P = 0.22) and all two-way interactions were far from being significant (P > 0.1 in all cases).

**Figure 3.**
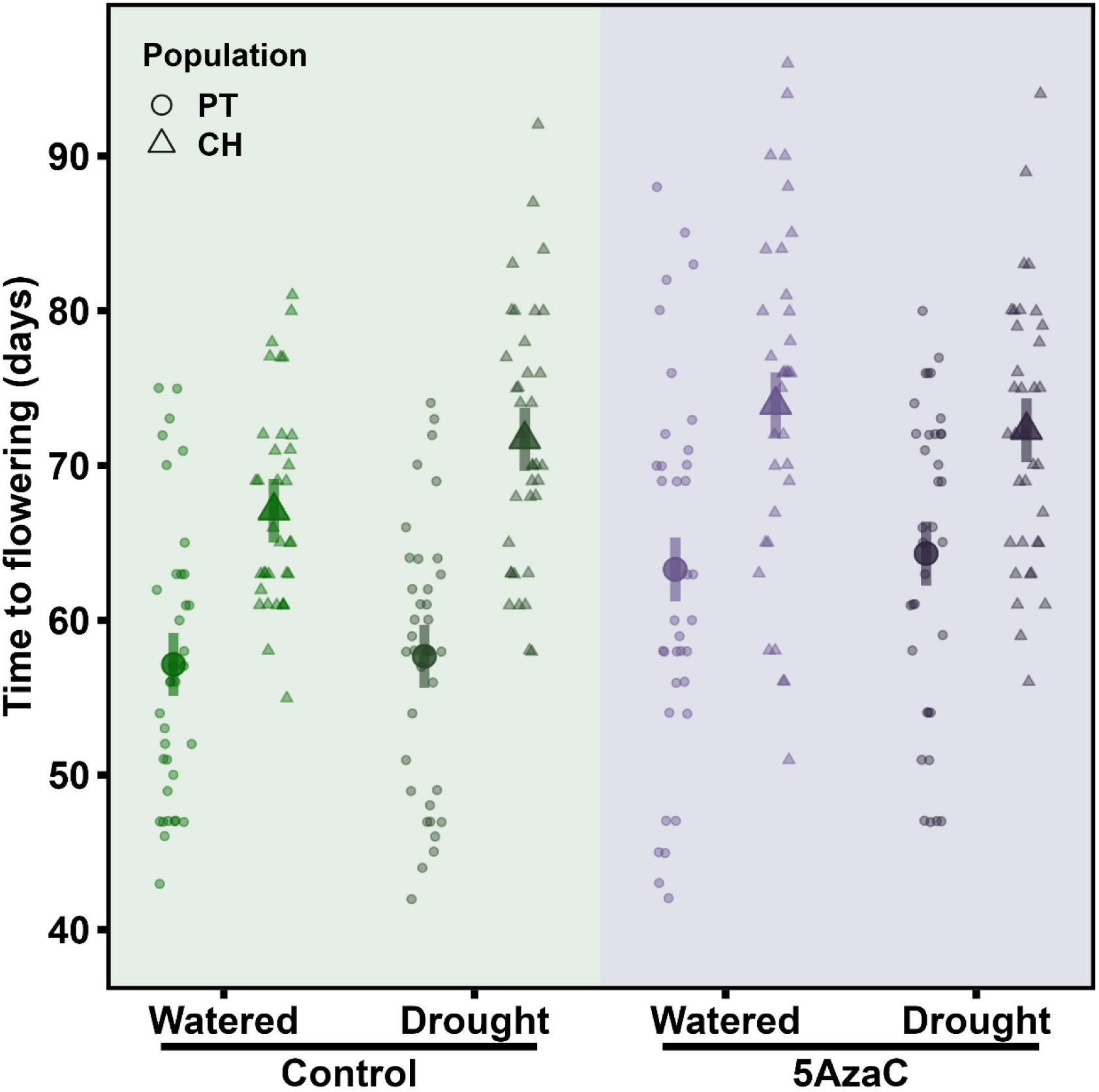
Effects of seed demethylation and recurrent drought treatments in time required to start flowering of *Erodium cicutarium* plants grown under controlled greenhouse conditions. Small circles and triangles denote individuals from PT and CH population, respectively. A darker color was used for individuals experiencing recurrent drought. Big symbols and bars represent leas-squared means and their 95% confidence level, respectively.

As regards plant growth, the production of new leaves changed with time showing a linear positive relationship (effect size estimate: 8.21 ± 0.22) and a negative quadratic term (effect size estimate: -6.17 ± 0.22) during the first nine weeks of plant growth. Demethylation treatment interacted significantly with the quadratic term (effect size estimate: -3.08 ± 0.42; Supp. Mat. Fig. S3A) indicating that control plants, on average, tended to produce less new leaves at bolting time (W9) whereas the demethylated plants recorded a more stable leaf production at that stage.

The number of new inflorescences produced every four weeks all through the reproductive period of the plants (W9-W17) showed a linear positive relationship with time (effect size estimate: 17.92 ± 0.47). Demethylation treatment interacted significantly with time (effect size estimate: -1.89 ± 0.88; Supp. Mat. Fig. S3B) indicating that control plants experienced a reduction in the number of inflorescences produced at late stage whereas demethylated plants did not change the rate of inflorescence production during the second period analyzed. The effect of recurrent drought did not interact in the production rate of leaves or inflorescences.

### Final individual size and fitness

Only recurrent drought had a significant and negative effect on above-ground plant biomass (ꭓ^2^ = 20, df = 1, P < 0.0001). Plants from the two provenances reached similar final aerial size (3.89 ± 0.12 g and 3.74 ± 0.14 g for CH and PT plants, respectively; ꭓ^2^ = 0.66, df = 1, P = 0.42). Demethylation and all interactions between study factors were far from being statistically significant (P > 0.4).

Root biomass was always smaller than aerial biomass, and the two traits were positively correlated across well-watered plants (rs = 0.30, P = 0.037, N = 48) but not in those experiencing recurrent drought (rs = 0.03, P = 0.84, N = 48). Recurrent drought had a significant and negative effect on root biomass (ꭓ^2^ = 33.5, df = 1, P < 0.0001). Moreover, root biomass was larger for CH plants (1.16 ± 0.06 g) than for PT plants (0.91 ± 0.06 g; ꭓ^2^ = 7.9, df = 1, P = 0.005). Demethylation and all interactions between study factors were far from being statistically significant in explaining individual variation in final root biomass (P > 0.4). Finally, PT plants exhibited a broad range of shoot:root biomass ratio (1.7 – 7.8; excluding an outlier = 10.9) that increased in water stressed plants (ꭓ^2^ =10.1, df = 1, P < 0.0015), whereas the biomass ratio varied less for CH plants (2.1 – 6.1; excluding an outlier = 10.4) and it did not change under stress (ꭓ^2^ = 0.21, df = 1, P = 0.6).

As regards reproductive output, the total number of inflorescences produced per plant by the end of the experiment averaged 42.7 and ranged between 15 and 94 (three individual outliers excluded produced 118, 121 and 122). Recurrent drought had a significant and negative effect on total inflorescence production (ꭓ^2^ = 13.6, df = 1, P = 0.0002). The effect of seed demethylation was also statistically significant (ꭓ^2^ = 4.5, df = 1, P = 0.033) with plants grown from 5azaC treated seeds producing on average 3.8 ± 2.6 fewer inflorescences than their untreated relatives (Fig. 4A). Furthermore, plants from CH produced on average 5.0 ± 3.2 fewer inflorescences than PT plants (ꭓ^2^ = 5.9, df = 1, P = 0.015). All interactions between study factors were far from being statistically significant (P > 0.19).

**Figure 4.**
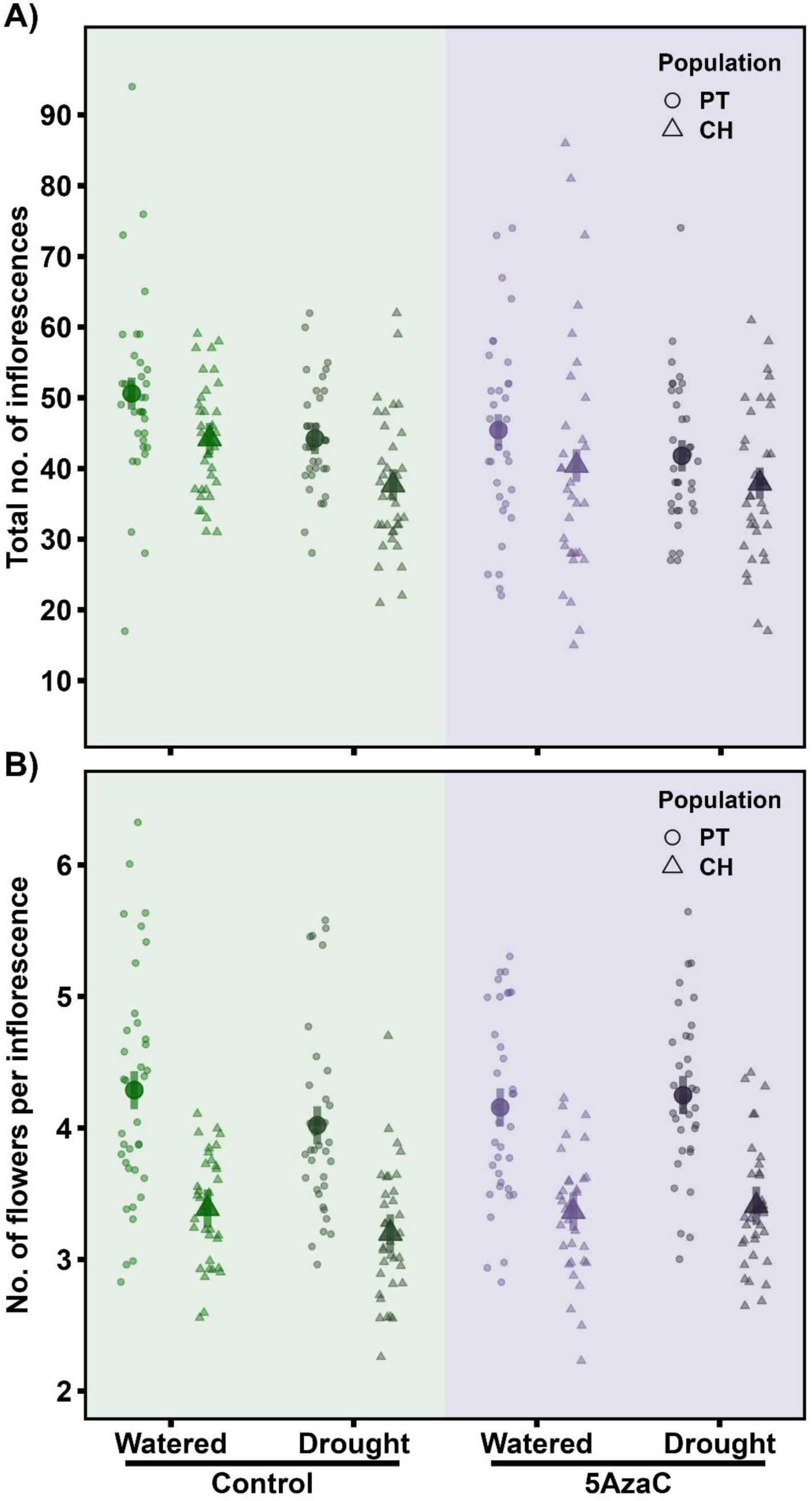
Effects of seed demethylation and recurrent drought treatments in flower investment of *Erodium cicutarium* plants. A) Total number of inflorescences produced. B) Average number of flowers produced per inflorescence. Small circles and triangles denote individuals from PT and CH population, respectively. A darker color was used for individuals experiencing recurrent drought. Big symbols and bars represent leas-squared means and their 95% confidence level, respectively.

The investment on reproductive output also depended on the number of flowers produced per inflorescence that ranged between 1 and 13. This parameter was again significantly lower in CH plants (4.18 ± 0.13 and 3.34 ± 0.13, for PT and CH plants, respectively; ꭓ^2^ = 21.0, df = 1, P < 0.0001; Fig. 4B) indicating that PT plants invested more in both reproductive parameters. For the first time, we found here a significant interaction between demethylation and drought (ꭓ^2^ = 7.5, df = 1, P = 0.006) indicating that recurrent drought induced a higher production of flowers per inflorescence only when plants were previously exposed to the 5azaC treatment (Fig. 4B), i.e. when the number of inflorescences produced was lower. All the other interactions were far from being significant (P > 0.5).

Plants grown from 5azaC treated seeds had lower reproductive success (Fig. 5). The effect of seed demethylation on fruit set probability was negative and statistically significant (ꭓ^2^ = 12.08, df = 1, P = 0.0005) and a significant interaction with provenance (ꭓ^2^ = 6.5, df = 1, P = 0.01) indicated that the effect of 5azaC was stronger in CH plants. Further, fruiting success was higher in plants that experienced recurrent drought although the effect was marginally significant (ꭓ^2^ = 3.1, df = 1, P = 0.08). Neither provenance nor any other interaction were statistically significant (P > 0.28). Similar results were found for the proportion of seeds set (Fig. 5B). The effect of seed demethylation was negative and statistically significant (ꭓ^2^ = 5.1, df = 1, P = 0.02) and a significant interaction with provenance (ꭓ^2^ = 7.3, df = 1, P = 0.007) indicated that the effect of 5azaC was stronger in CH plants. Recurrent drought increased the probability of producing seeds per ovule (ꭓ^2^ = 5.7, df = 1, P = 0.017). Neither provenance nor any other interaction were statistically significant (P > 0.19).

**Figure 5.**
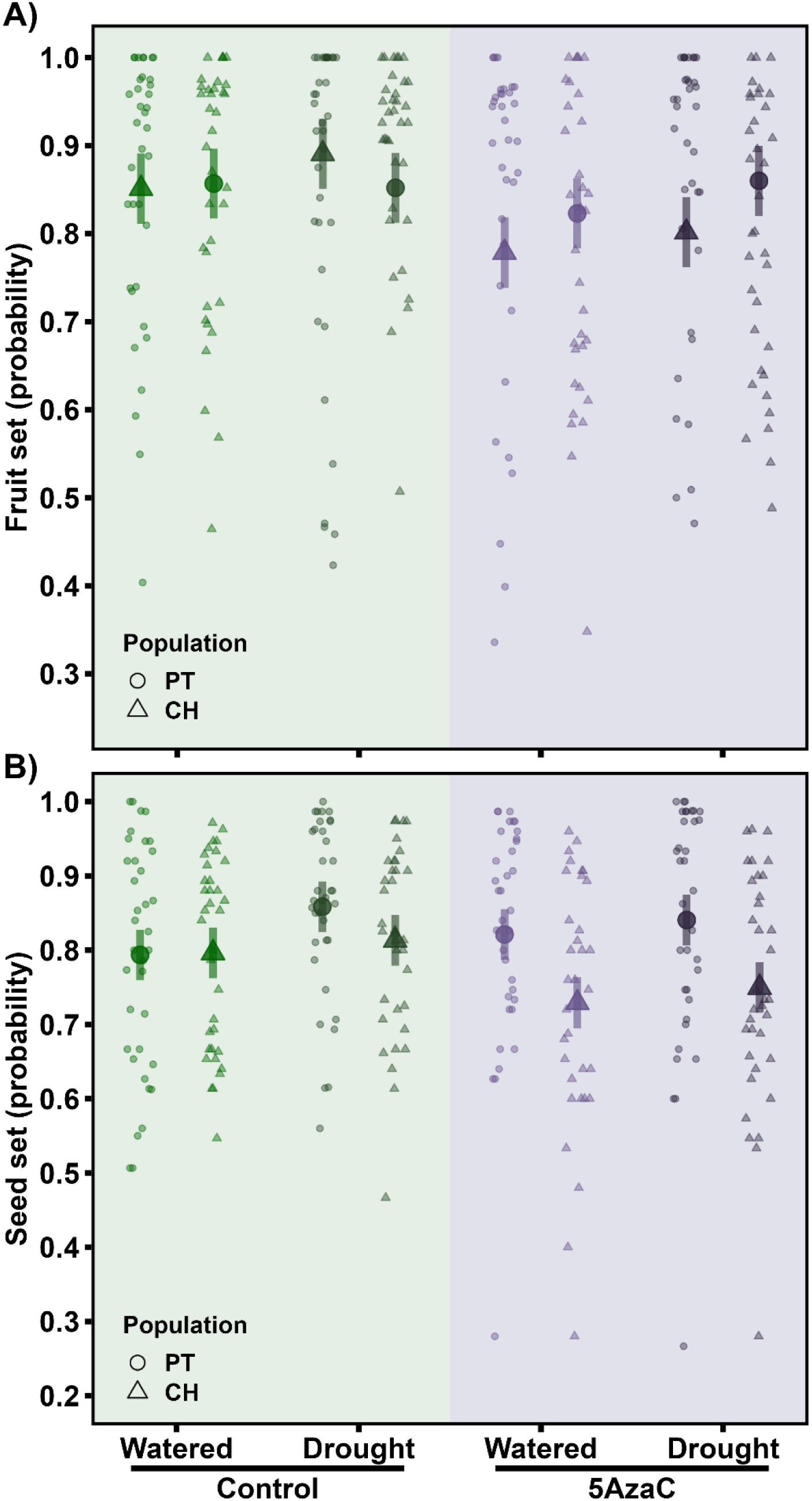
Effects of seed demethylation and recurrent drought treatments in reproductive success of *Erodium cicutarium* plants. A) Average proportion of flowers setting fruits. B) Average proportion of ovules setting seeds within fruits. Note this species is autonomous for producing fruits and has a fixed number of five ovules per fruit. Small circles and triangles denote individuals from PT and CH population, respectively. A darker color was used for individuals experiencing recurrent drought. Big symbols and bars represent leas-squared means and their 95% confidence level, respectively.

### Phenotypic variability associated to actual variation in global DNA methylation

Finally, we analyzed the continuous linear and quadratic relationships between phenotypic traits and global DNA methylation in leaf and root tissues. Results are summarized in Table 2. We found global cytosine methylation in leaf DNA was better related to developmental ontogeny-related traits. The most significant results indicated that plants with lower global cytosine methylation in leaf DNA produced shorter first leaves, had less leaves at early stage (11 days), and required some more time to start flowering, compared to those with intermediate and high global leaf DNA methylation percentage.

**Table 2.** Summary of the results of the linear mixed models conducted to analyze the continuous relationships between phenotypic traits and global DNA methylation in A) leaf tissue and B) root tissue. Both linear and quadratic relationships were tested (N = 96 individuals), the quadratic term is only shown if model fit improved when included. Only phenotypic traits showing some close to significant relationship (P < 0.10) are included, significant relationship are highlighted in bold. All models included maternal family as a random effect.

| Response trait | Linear effect size | s.e. | Chi-squared | P | Quadratic effect size | s.e. | Chi-squared | P |
| --- | --- | --- | --- | --- | --- | --- | --- | --- |
| A) Leaf DNA |  |  |  |  |  |  |  |  |
| 1<sup>st</sup> leaf length | 5.95 | 1.29 | 21.17 | **<0.0001** |  |  |  |  |
| No. leaves 1 | 0.24 | 0.08 | 9.19 | **0.002** |  |  |  |  |
| No. leaves 2 | 9.36 | 5.08 | 3.38 | 0.07 | -0.22 | 0.12 | 3.42 | 0.06 |
| Flowering time | -16.9 | 5.4 | 9.51 | **0.002** | 0.37 | 0.13 | 7.68 | **0.006** |
| B) Root DNA |  |  |  |  |  |  |  |  |
| No. leaves 1 | -1.67 | 0.87 | 3.55 | 0.06 | 0.04 | 0.02 | 3.68 | 0.06 |
| No. leaves 2 | 0.62 | 0.27 | 5.25 | **0.02** |  |  |  |  |
| No. leaves 3 | 0.71 | 0.23 | 9.89 | **0.002** |  |  |  |  |
| Above ground biomass | 0.07 | 0.04 | 2.94 | 0.09 |  |  |  |  |
| Below ground biomass | 0.08 | 0.01 | 60.28 | **<0.0001** |  |  |  |  |

Variation in global cytosine methylation in root DNA was linearly and positively related to the number of leaves plants had at mid and late vegetative stages, and more significantly to final root mass (Table 2).

## Discussion

DNA cytosine methylation is a dynamic epigenomic feature that in plants can change in response to developmental and environmental cues (Finnegan, 2010; Brautigam and Cronk, 2018). Analyzing how these changes can be interrelated is key to better understanding epigenetic contributions in plant response to stress, that could lead or not to phenotypic plasticity and priming depending when it happens during their life-cycle (Brautigam and Cronk, 2018; Cooper and Ton, 2022). The use of experimental demethylation at seed stage enlarge variance in global DNA methylation and may help to explore the epigenetic and phenotypic outcome of stress exposure at both above and below-ground. In this study, we found that an initial short treatment of *E. cicutarium* seeds with 5azaC was still detectable in roots and leaves as a lower global cytosine methylation in DNA when plants reach reproductive stage, supporting the utility of this protocol to alter individual methylation profiles (see Balao *et al.,* 2023 for a deeper molecular study). The ontogenetic phenotypic characterization was able to unveil a delay in development and growth after seed demethylation, whilst the static final phenotypic characterization showed a reduction in size and inflorescence production after recurrent drought, with partial compensation of seed output by increased autonomous reproductive success. In the following paragraphs, we will discuss in more detail the consequences of seed demethylation and recurrent drought in terms of global DNA methylation, plant development and individual final size and fitness-related traits.

### A dynamic analysis of global DNA methylation

Global DNA methylation is known to increase in leaf tissue from seedling to adult age in *Arabidopsis thaliana* (Baubec *et al.,* 2009) and a few other species (e.g., Perrin *et al.,* 2019; Alves *et al.,* 2020). Furthermore, DNA methylation can change among tissues (Messeguer *et al.,* 1991; Demeulemeester *et al.,* 1999; Seymour *et al.,* 2014; Lloyd and Lister, 2022) and its variation across growth stages (or dates) may change differently on different tissues depending on environmental conditions (Demeulemeester *et al.,* 1999; Seymour *et al.,* 2014; Lloyd and Lister, 2022). In well-watered -control-plants of *E. cicutarium*, global DNA methylation varied between leaf and root tissues and increased from seedling to adult stage, difference with age being stronger in root tissue (from 13.0 % to 22.8 %) than in leaf tissue (from 17.7 % to 20.9 %). Therefore, under benign conditions, DNA was more methylated in leaves than roots at seedling stage (Alonso *et al.,* 2017) whereas the opposite was true at the adult stage, as reported here.

Cytosine methylation in DNA obtained from adult leaves and roots was positively correlated in well-watered plants. However, in those individuals experiencing recurrent drought along their lifetime, global DNA methylation in the two tissues was decoupled, suggesting that epigenetic processes above- and below-ground could be to some extent self-regulated and independent from each other (see also Abid *et al.,* 2017 for evidence in seedlings of *Vicia faba*). The absence of correlation most likely reflects that recurrent water stress consistently and largely reduced methylation in adult root DNA, whereas in leaves reduction in DNA methylation along development was milder and unrelated to water regime. At molecular level, the complexity of interaction effects between seed demethylation and recurrent drought was revealed as eight patterns of methylation change in cytosines located in different fragments of leaf DNA (Balao *et al.,* 2023). In many cases, 5azaC and drought induced methylation changes on different cytosines and the combination of the two treatments re-established methylation level to that found in control plants (see Balao *et al.,* 2023 for further details). Ongoing analyses on the molecular changes occurring at root DNA should shed light on the extent to which epigenetic regulation in response to seed demethylation and other external factors such as drought can differ between above and below-ground tissues (Lloyd and Lister, 2022).

### Developmental and fitness consequences of seed treatment with 5azaC

The short seed exposure to 5azaC applied here did not affect early seedling mortality. It slowed individual growth, delayed flowering and leaf production, and had a negative impact on fitness by reducing both inflorescence production and reproductive success. Our results therefore suggested that the epigenetic effects can desynchronize plant growth, flowering and senescence among individual plants in both favourable and adverse environments (see Herrera *et al*., 2019 for similar effect within plants). Early studies on the effects of 5azaC highlighted its ability to simulate vernalisation, establishing links between DNA methylation and flowering transition (Fieldes and Amyot 1999; Kondo *et al.,* 2007). At population level, both flowering time and flowering synchrony are able to impact on fitness particularly in annual plants and temperate climates (Munguía-Rosas *et al.,* 2011). An epigenetic regulation of flowering time could be particularly beneficial in unpredictable environments and in response to climate change (Franks & Hoffman, 2012). For instance, in Mediterranean mountains shorter growing periods associated to earlier and hotter dry summers could favour early blooming individuals in some years but not in others (Giménez-Benavides *et al.,* 2010) and, thus, reversible epigenetic changes could be particularly advantageous. Future studies combining the analysis of methylome and transcriptome effects of 5azaC in different plant species or ecotypes could help to better characterize the pathways linking DNA methylation and flowering (see e.g., Wang *et al.,* 2017 for the analysis of flowering induction by ethylene in pineapple).

### Changes in final size and fitness induced by recurrent drought and the interaction with 5azaC

*Erodium cicutarium* plants could not fully compensate the negative impact of recurrent water shortage and attained lower individual size and inflorescence production without significant effects on phenology or growth-related traits. Interestingly, stressed plants were able to partially compensate their reproductive output. First, recurrent drought induced a higher production of flowers per inflorescence only if plants had been previously exposed to the 5azaC treatment. As regards the seed production of this autonomous plant, fruit set and seed set probabilities increased under recurrent drought whereas they were negatively impacted by seed demethylation. Such findings suggested that allocation of resources to reproduction could be to some extent upregulated by stress and mediated by DNA methylation, likely through their combined effects on hormones’ metabolism (Fleta-Soriano and Munné-Bosch, 2016; Avramova, 2019; Ezura *et al.,* 2023).

### Heterogeneous response between seed provenances: from experimental nuisance to ecological relevance

The delay in flowering and growth induced by seed demethylation and decreased final reproductive output induced by recurrent drought varied in magnitude between the two seed provenances analysed despite the short distance between source populations. Such unexpected result suggested that not only intraspecific variability associated to broad geographical scales and climatic gradients may determine the magnitude of plant phenotypic plasticity mediated by DNA methylation changes (e.g., Bossdorf *et al*., 2010; Münzbergová *et al*., 2019; Sammarco *et al*., 2021). Short-distance environmental heterogeneity emerged as another fundamental scale leading to intraspecific epigenetic variability (e.g., Herrera & Bazaga, 2016; Herrera *et al*., 2017; Valverde *et al*., 2024) whose relevance for plant adaptation to stress deserves further analysis. First, we showed that the magnitude of reduction in DNA methylation of leaves and roots after seed treatment with 5azaC was not homogeneous between provenances (see e.g., Troyee et al., 2022 for heterogenous response between two ecotypes of *Thlaspi arvense*, and Browne *et al*., 2020 for heterogenous response in gene transcription between families of *Quercus lobata*). As a practical consequence, the predicted interaction between our two fixed factors was somehow weakened by enlarged within class variance and the statistical power for our 3-way interaction model was reduced. Second, we found that individuals from PT flowered earlier and, according to our expectations for the scape-avoidance strategy, were less affected by drought, being able to produce more flowers per inflorescence and higher fruit- and seed-set probabilities. Interestingly, they were also the individuals whose DNA methylation in leaves and roots was decoupled by drought, suggesting that a more responsive methylome could to some extent blur the seed treatment effect and contribute to improve fitness under stress.

## Conclusions

By looking into changes in global DNA methylation in leaves and roots of *E. cicutarium* we have investigated the heterogeneous phenotypic response to water availability within and across individuals. Within individuals, we documented some independence for epigenetic regulation above- and below-ground along plant development, and how drought elicited a stronger phenotypic and molecular change in roots. Across individuals, we found intraspecific variation in the ability to change plant methylome in response to drought in this annual plant, supporting that an escape-avoidance strategy could be mediated by changes in plant methylome.

## Acknowledgements

We thank Maciej Barczyk, Laura Cabral, Esmeralda López, Daisy Johnson, Pablo Martín, Alejandro Mira, Ricardo Pérez, Elena Villa and Noelia Zarza, for assistance in field, greenhouse and laboratory work, and Javier Puy for suggestions on a draft version of the manuscript. Financial support was provided by grants EPIECOL-CGL2013-43352-P, EPIENDEM-CGL2016-76605-P, EPINTER-PID2019-104365GB-I00 and DISTEPIC-PID2022-141530NB-C22 (Ministerio de Ciencia, e Innovación, Spanish Government).

## Author contributions

All authors designed the research; C.A. and M.M. conducted greenhouse experiment, data collection and data analyses; C.M.H. and C.A. provided economic resources; C.A. led the writing; M.M. prepared all figures; all authors contributed to refining the manuscript.

## Figures for Supplementary Materials

**Supp Mat Fig S1.**
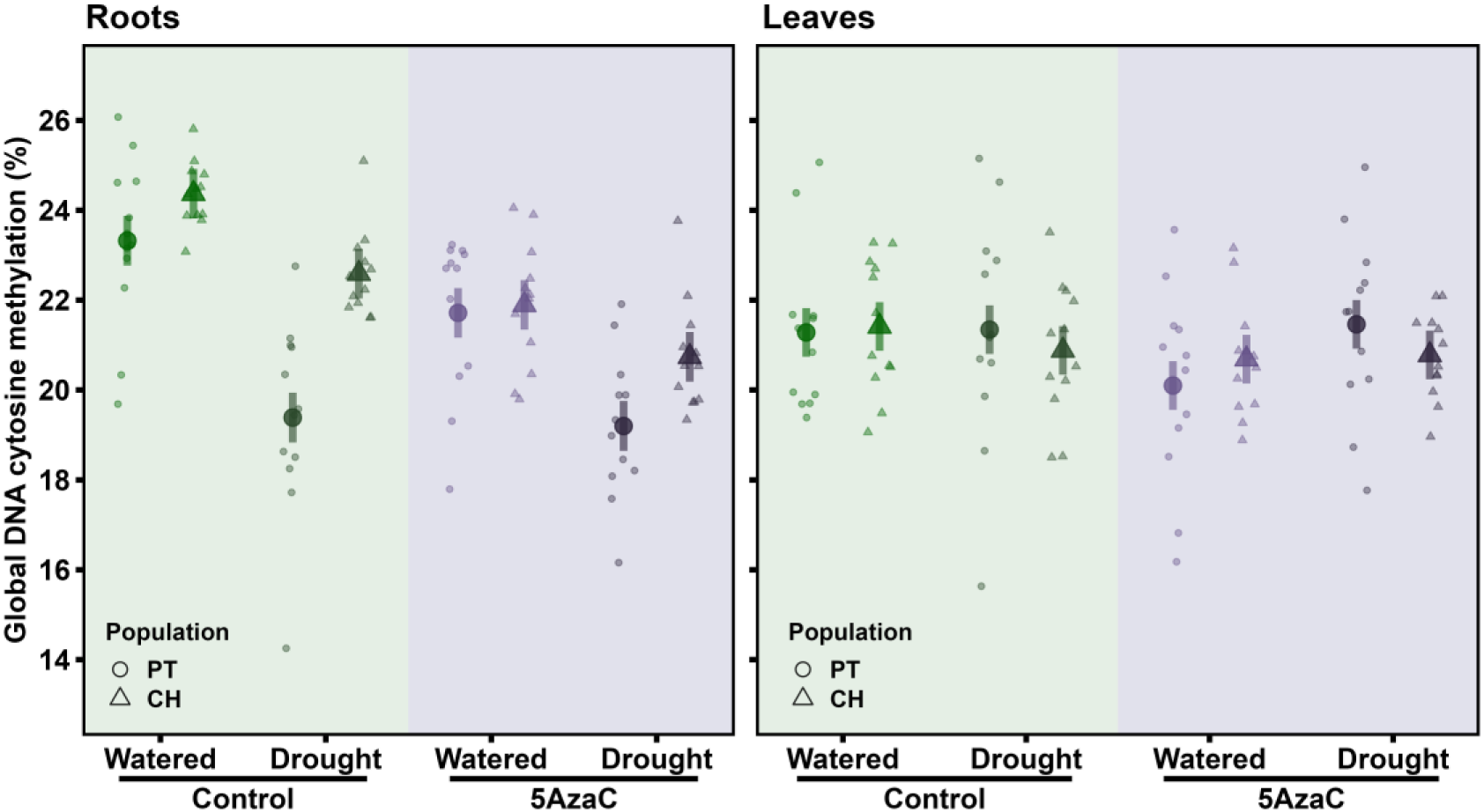
Effects of seed demethylation and recurrent drought treatments in global DNA cytosine methylation percentage of *Erodium cicutarium* plants. A) DNA obtained from root tissue. B) DNA obtained from leaf tissue. Small circles and triangles denote individuals from PT and CH population, respectively. A darker color was used for individuals experiencing recurrent drought. Big symbols and bars represent leas-squared means and their 95% confidence level, respectively.

**Supp Mat Fig S2.**
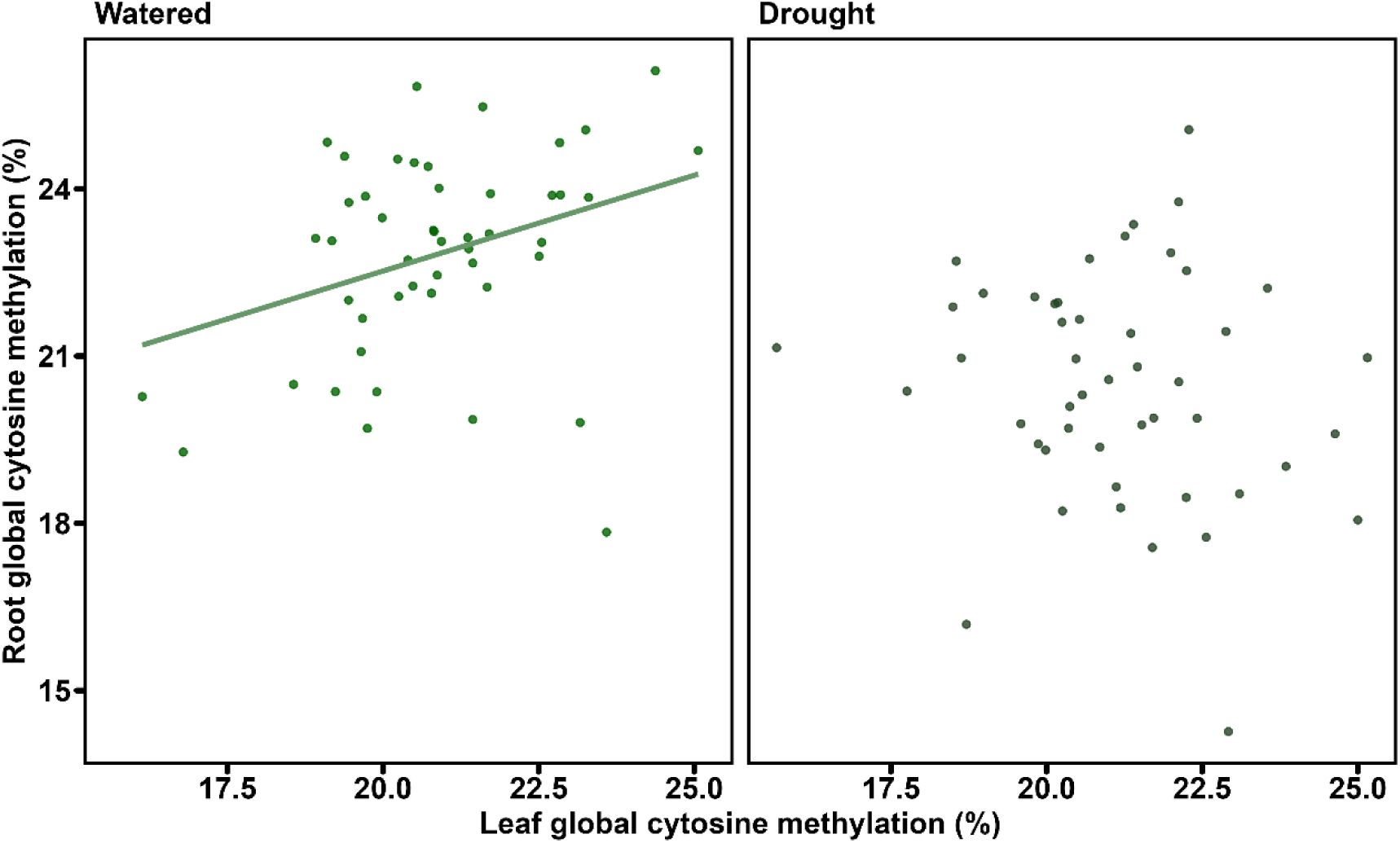
Graphical representation of the relationship between global percent of methylated cytosines in DNA obtained from leaves and roots of individual adult plants according to the level of water availability experienced along their lifetime. Each dot represents an individual and colors were used to distinguish individuals that were regularly watered (green) or experienced recurrent drought (grey). The regression line is represented only when significant.

**Supp Mat Fig S3.**
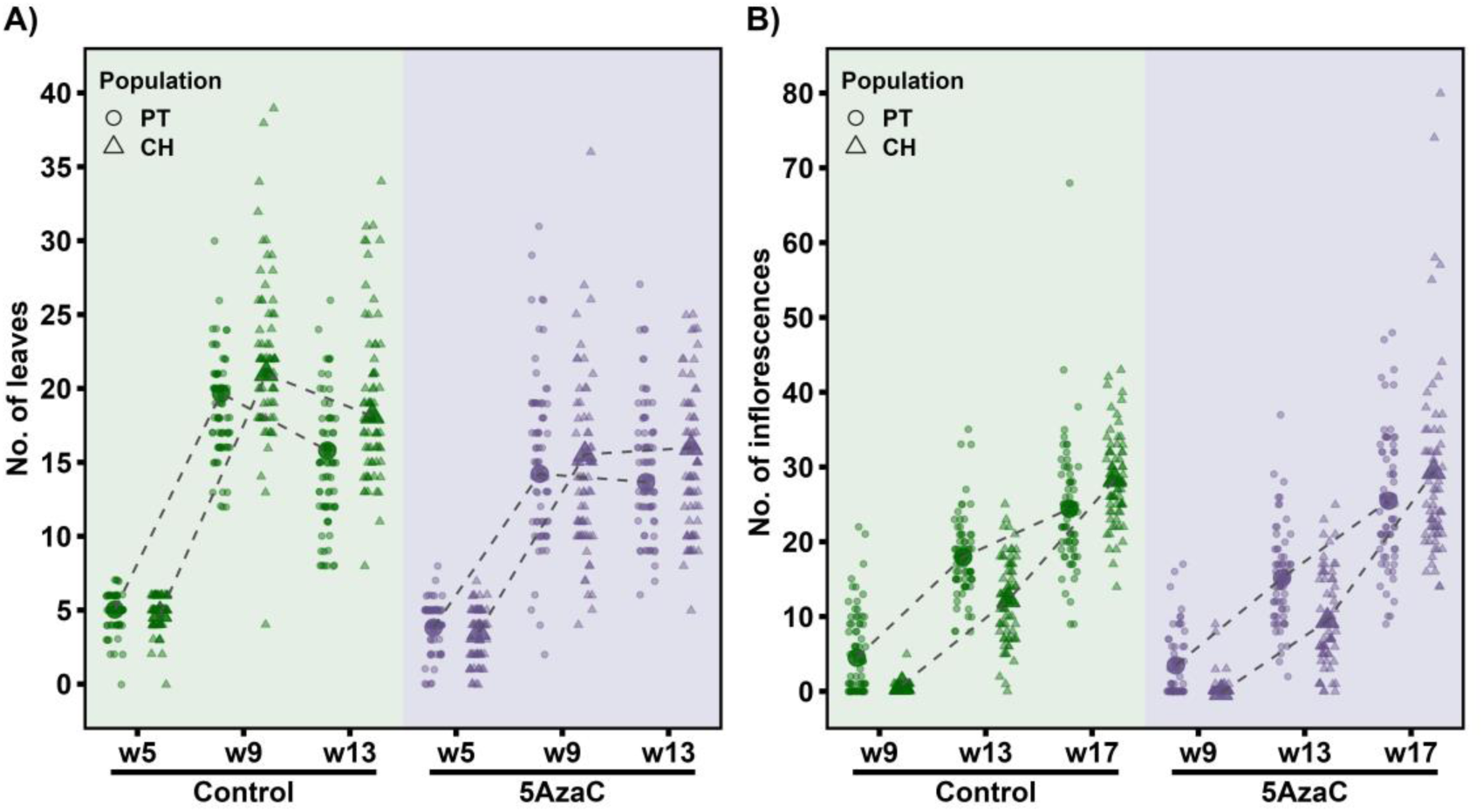
Effect of seed demethylation treatment in developmental phenology of *Erodium cicutarium* plants A) Number of full-grown leaves per individual recorded on different weeks (W1-W9). B) Number of inflorescences produced on different weeks (W9-W17). Small circles and triangles denote individuals from PT and CH population, respectively. Big symbols and bars represent leas-squared means and their 95% confidence level, respectively.

